# Magnetic Temporal Interference for Noninvasive Focal Brain Stimulation

**DOI:** 10.1101/2022.01.10.475762

**Authors:** Adam Khalifa, Seyed Mahdi Abrishami, Mohsen Zaeimbashi, Alexander D Tang, Brian Coughlin, Jennifer Rodger, Nian X. Sun, Sydney S. Cash

## Abstract

Non-invasive stimulation of deep brain regions has been a major goal for neuroscience and neuromodulation in the past three decades. Transcranial magnetic stimulation (TMS), for instance, cannot target deep regions in the brain without activating the overlying tissues and has a poor spatial resolution. In this manuscript, we propose a new concept that relies on the temporal interference of two high-frequency magnetic fields generated by two electromagnetic solenoids. To illustrate the concept, custom solenoids were fabricated and optimized to generate temporal interfering electric fields for rodent brain stimulation. C-Fos expression was used to track neuronal activation. C-Fos expression was not present in regions impacted by only one high-frequency magnetic field indicating ineffective recruitment of neural activity in non-target regions. In contrast, regions impacted by two fields that interfere to create a low-frequency envelope display a strong increase in c-Fos expression. Therefore, this magnetic temporal interference solenoid-based system provides a framework to perform further stimulation studies that would investigate the advantages it could bring over conventional TMS systems.

## I. Introduction

The clinical use of transcranial magnetic stimulation has been a prominent achievement in the field of neuroscience in the past two decades [1]–[3]. The electric field (E-field) generated by the TMS coils can be categorized as subthreshold or suprathreshold. Suprathreshold E-fields are large enough to directly elicit action potentials in neurons while subthreshold E-fields generated by repetitive transcranial magnetic stimulation (rTMS) can induce neural plasticity [4]. rTMS, as an FDA-approved technique, has provided remarkable relief to patients with depression, migraine, obsessive-compulsive disorder (OCD), and other debilitating disorders [2], [5], [6]. As most noninvasive brain stimulation technologies, TMS suffers from two fundamental limitations that the scientific community has been unable to solve during the past two decades:

1. TMS is mainly limited to superficial cortical brain sites as the electric field is always maximal closest to the stimulating coil. It generally reaches no more than 3 cm into the brain due to the rapid spatial attenuation of the magnetic and electric fields generated by the coil [7], [8]. This is unfortunate as it has been demonstrated that the regions involved in most neurological and neuropsychiatric disorders are also located in non-superficial brain areas and accessing these deep structures could be an efficient treatment for patients with depression, OCD, Parkinson’s disease, essential tremor, and dystonia. Novel coils such as the H-coil has the ability of deep brain stimulation but is unable to do so without activating the overlying tissues [9], [10].
2. Despite recent efforts that have led to the development of new coil designs (new shape and size) to improve the focality and stimulation depth, TMS still shows poor spatial resolution [11]–[14]. This is caused by the large size of conventional TMS coils, which in addition to target sites, also stimulate neighboring brain regions and lead to side effects such as headache, twitching of facial muscles, or lightheadedness [15].

In this research, to improve stimulation depth and focality, we propose a novel TMS technique which we call magnetic temporal interference (MTI). MTI relies on two magnetic solenoids that are driven at two slightly different high frequencies (e.g., 10.5 kHz and 11 kHz). Non-target regions of the brain are impacted only by one high-frequency magnetic field (e.g., 10.5 kHz or 11 kHz) which alone is ineffective at recruiting neural activity, while on-target regions are impacted by two fields that interfere to create a low-frequency envelope (e.g., 500 Hz) as shown in Fig. 1. The constructive interference hot spot can be precisely controlled by adjusting the solenoid placement and delivered current. In this research, we will focus on repetitive magnetic temporal interference (rMTI). Analogous to transcranial alternating current stimulation (tACS), rMTI generates continuous oscillating fields, as opposed to rTMS which delivers brief bursts. Prior neuromodulation studies that have relied on temporal interference have shown that neurons can be activated by the low-frequency envelope [16], [17]. It was demonstrated in animal models with focused ultrasound stimulation [16] and tACS [17] but has yet to be experimentally demonstrated using magnetic stimulation. Therefore, the goal of this research is to: i) construct solenoids optimized for high frequency (kHz) magnetic stimulation, and ii) show that rMTI stimulation induces focal stimulation using a rodent model.

**Fig. 1:**
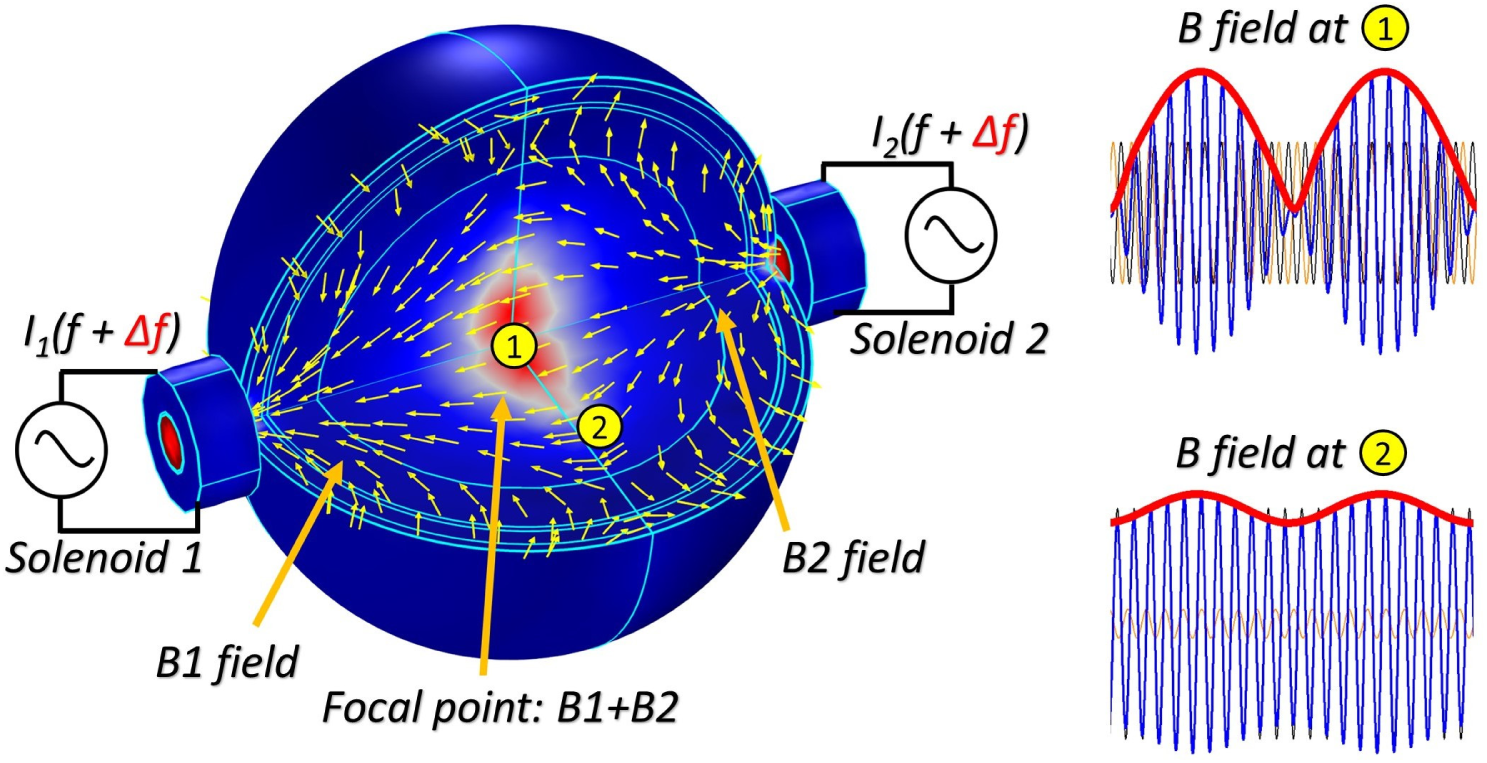
Simulation results showing the concept of rMTI on a 20 cm diameter brain model which includes different layers (i.e., skin, skull, CSF, grey matter, and white matter). Solenoid 1 and 2 generate a high frequency sinusoidal wave which interfere at the center of the brain to create a low frequency sinusoidal wave.

The rest of the manuscript is organized as follows. Section II describes the methods used for simulation, design, and fabrication of MTI solenoids. Section III presents the simulation and characterization results and demonstrates the MTI stimulation proof-of-concept which was done by quantifying immediate early gene expression (c-Fos) as a measure of neural activity. Finally, Section IV discusses the results and raises questions about the stimulation mechanism – how does the rMTI E-field influence the transmembrane potential of neurons to induce activity or plasticity?

## II. Method

### A. Solenoid design and characterization setup

One of the most important parameters to select when designing resonating solenoids for rMTI stimulation is the carrier frequency and the beating frequency (i.e., the frequency offset between the carrier frequencies). We know that the electric field linearly increases with frequency based on Faraday’s Law of electromagnetic induction (i.e., the induced electric field is linearly proportional to the rate of change of the magnetic field over time). We also know, based on past works [18]–[20] that looked at the link between neural response and sinusoidal electric fields, that the sensitivity of transmembrane potentials to sinusoidal electric fields decreases with increasing stimulation frequency. This could be due to the relatively small neural membrane time constant which prevents the neural electrical activity from following very high frequency oscillating electric fields. Therefore, there exists a “sweet spot” for the carrier frequency that enables the most effective rMTI stimulation. We chose 10.5 kHz and 11 kHz as the carrier frequencies based on our understanding of prior work that characterized high-frequency electrical stimulation [19], [21]. However, we would like to note that this is most likely not the optimal value since simulations of neural models often lack accuracy and do not take into account the complexity of the brain. The beat frequency was chosen to be 500 Hz. We decided to pick a time period that is shorter than the membrane time constant so that after a certain number of cycles, the induced membrane fields could potentially reach the neuron firing threshold. In other words, although one single cycle will generate subthreshold E-fields, the entire waveform (which is composed of 10 cycles) has the potential for suprathreshold stimulation.

If the coil or solenoid is not miniaturized sufficiently then it can result in the stimulation of the entire brain of the rodent [22]–[24]. Therefore, a rat-specific stimulating solenoid needed to be designed for this research. The MTI solenoid parameters (e.g., diameter, length, number of turns, and magnetic core) and their location on the brain are optimized based on 2 criteria:

1. The produced magnetic/electric fields by two solenoids can focally interfere at targeted sites.
2. The quality factor 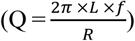 of the solenoid is maximized. *L* is the effective inductance, *R* is the wire effective resistance, and *f* is the operating frequency. A larger Q leads to a larger generated electric field, which is the most important parameter for magnetic stimulation. In addition, a large Q could also signify that the resistance is small which would allow the solenoid to handle larger currents. When optimizing solenoids, it is important to remember that there exists a tradeoff between focality and electric field strength. For instance, the Q can be increased by using larger diameter wires or increasing the number of turns, but this would result in an increase in solenoid diameter which would decrease focality.

Using a solenoid instead of a coil (like in commercial TMS systems) allows us to use magnetic cores to boost the generated magnetic/electric fields. The magnetic core is made of annealed carbonyl iron powder material (Mix 1, Micrometals, U.S.A). The core can operate at frequencies up to 1 MHz (much larger than our operating frequency). Powder cores were chosen over ferrite cores for their larger saturation flux density (Bsat). Ferrite cores were initially used as they provided larger relative permeabilities, however, they led to less efficient solenoids due to lower core saturation magnetization. We would like to note that even with powder cores, the operating current during rMTI stimulation (i.e., 150 A) is still much higher than the saturation current which is estimated to be 13A. We decided to taper the magnetic core using a Dremel tool to channel the magnetic flux to a smaller target within the brain. This technique allowed us to avoid the need to further reduce the solenoid diameter which would have increased the generated heat.

We used a search coil made in-house to measure the magnetic field density along the z-axis (B_z_-field) generated by the solenoid. The search coil had 4 turns and an inner diameter of 0.8 mm. The number of turns and diameter were reduced as much as possible to increase resolution. The recorded voltage, which was converted to B_z_-field, had to be measurable. When small B_z_-fields were applied, we found that search coils with less than 4 turns led to small recorded voltages that were below the noise floor of the oscilloscope.

Two setups were used to characterize the solenoids with the objective of comparing measured B_z_-field results with simulation B_z_-field results to evaluate the accuracy of our COMSOL simulations. In the first setup (Fig. 2(a)) the search coil was placed 2 mm from a vertically aligned solenoid with a center-to-center alignment. The second setup is described in the next section. Stimulation parameters were controlled by a waveform generator (SDG2042X, Siglent, USA) which was connected to a power amplifier (Magnitude3600, Seismic Audio, USA). Setting the power amplifier in bridge mode allows it to output a maximum power of 3600 Watts RMS with 8 Ω connected load.

**Fig. 2:**
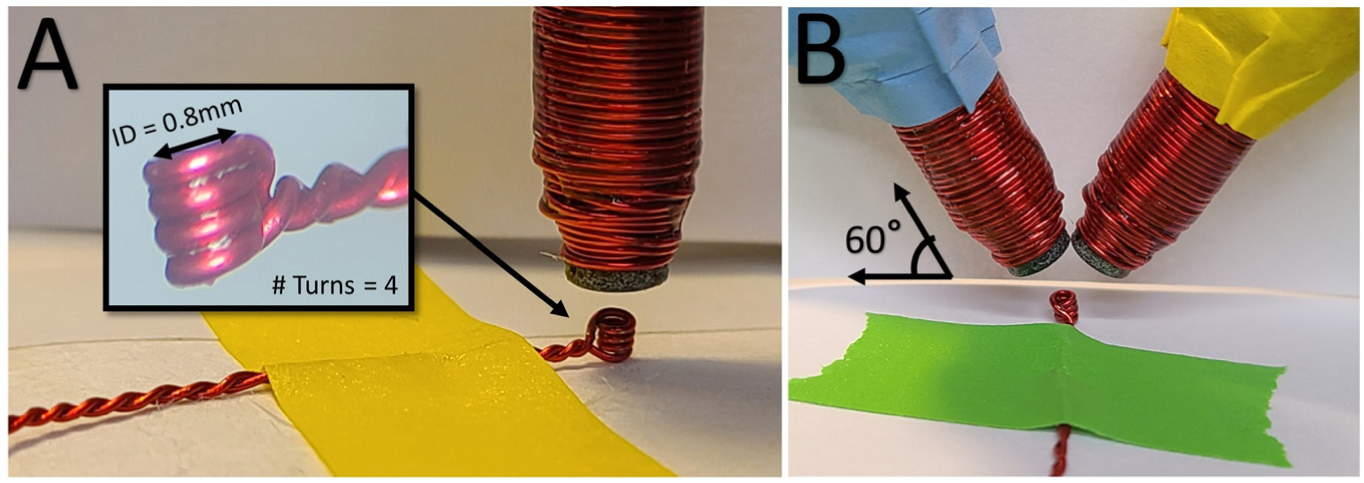
Photographs of the solenoid characterization setup. (A) Setup for measuring the B_z_-field of 1 solenoid with a close-up view of the search coil. (B) Setup for measuring the temporally interfering B_z_-field induced by 2 solenoids.

### B. MTI characterization setup

To create MTI, at minimum 2 solenoids need to be used simultaneously. The focal point where the two fields interfere can then be adjusted at any depth inside the brain by changing the location of the two solenoids or the value of the currents injected into the solenoids. Each solenoid was fixated onto a stereotaxic micropositioner which allowed for accurate and controlled positioning. Both solenoids were tilted such that they formed a 60° angle as shown in Fig. 2(b). The temporally interfering B_z_-field was measured using the search coil placed 2 mm from the solenoids. This distance was chosen since it roughly corresponds to the distance between the solenoid tip and the surface of the cortex. The current provided to each solenoid is then slightly adjusted to optimize the interference.

One of the major issues, when solenoids/coils are miniaturized for rodent magnetic stimulation, is the large amount of heat they generate. When the solenoid temperature increases, the Bsat of the magnetic core decreases, and the wire impedance increases. Most importantly, excessive heating in the solenoid can potentially harm the rodent. It is thus crucial to set a limit on the amount of heat generated by the solenoids, which results in a maximum allowable current amplitude supplied to the solenoids. We placed the solenoids at 0.5 mm above the scalp of the rodent and measured the solenoid and scalp temperatures using a very high-resolution thermal imaging camera (TIM 640, Micro-Epsilon, USA) during rMTI stimulation. A fan was blowing towards the solenoids to limit the increase in temperature.

### C. Finite Element Modeling

All simulations were done in COMSOL Multiphysics software. The characterization setups described above are replicated in COMSOL. The 1^st^ setup replicates the characterization measurement setup of the single vertically aligned solenoid. The core saturation is taken into consideration in the simulation results. The 2^nd^ setup replicates the characterization measurement setup of the two tilted solenoids. Here we looked at the spatial profile of the magnetic flux density along the z-axis and the beating envelope (E_env_). To quantify the intensity of the beating envelope caused by the rMTI we used the following formula:

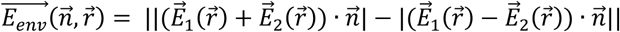

where 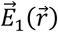 and 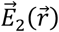 are the fields generated by the two solenoids at the location 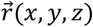, and 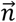 represents different directions (i.e, *x, y, z* axis). This allowed us to identify the hot spot(s) by plotting the spatial distribution of E_env_. The sinusoidal current applied to both solenoids was 150 A and runs only for 4 ms. Therefore, 2 full cycles of the 500 Hz envelope will be shown.

### D. In vivo experiment setup and c-Fos imaging

Although the end goal of rMTI stimulation is to be deployed into human clinical trials, as this manuscript introduces the concept, we relied on rodent models for its validation. The objective of the following proof-of-concept experiment was to show that rMTI stimulation is capable of increasing c-Fos levels (an indirect marker of neural activity), while stimulation with a single solenoid or two solenoids provided with the same signal does not lead to any detectable c-Fos increase.

Although past works have shown that the effects rTMS are better observed in awake animals [25], [26], we decided to conduct the animal experiment under light isoflurane anesthesia. Carefully aligning the solenoids for focal temporal interference is challenging when the rat is awake as rodents do not readily comply with motion restrictions. Furthermore, the noise emitted during MTI stimulation and the restraint of the rat would increase its stress level which could affect c-Fos expression [27]. Even though rMTI stimulation is meant to be used non-invasively (as in rTMS), in this first proof-of-concept we decided to place the solenoids directly above the skull for the following reasons: i) the landmarks on the skull make it easier to target a specific brain region, ii) solenoids are closer to the brain, and iii) it is easier to attach the search coil to the skull than to the scalp.

All research protocols were approved and monitored by the Massachusetts General Hospital (MGH) Institutional Animal Care and Use Committee (IACUC). Five male Sprague Dawley rats (Charles River Labs, MA, United States) were used in this study. Three rats received rMTI stimulation from 2 solenoids with different frequencies (10.5 kHz + 11 kHz), one rat received magnetic stimulation from 2 solenoids that received the same frequency signal (11 kHz + 11 kHz), and one rat received magnetic stimulation from 1 solenoid (11 kHz). The rMTI stimulation setup (i.e., waveform, instruments, configuration) is shown in Fig. 3(a). The rat was anesthetized with isoflurane throughout the surgery. After shaving the fur over the surgical site, the animal was immobilized on a stereotaxic frame (Fig. 3(b)), and the scalp was disinfected and numbed with lidocaine. A sagittal incision (up to 1.5 cm in length) was made in the skin over the skull. We then placed a mark on the skull to indicate the target site (i.e., from bregma, AP -3 mm, ML - 2.5 mm) and glued the search coil on the mark using cyanoacrylate (Fig. 3(c)). Having the search coil glued onto the skull before rMTI stimulation allowed us to adjust the currents provided to the solenoids to optimize the beating effect. After detaching the search coil, we placed the solenoids as close as possible to the skull (0.5-1 mm) at a 60° angle and decreased the isoflurane concentration down to 1% to mitigate the suppressive effects of anesthesia on neural activity. The power amplifier was then switched on and the solenoids stimulated the rat brain for 15 min using the same waveform parameters as those used during testbench tests (i.e., two sinusoids at 10.5 kHz and 11 kHz delivered for 20 ms, at 150 A and 1 Hz stimulation). During the first three experiments, we monitored the temperature of the solenoid, scalp, and skull. Finally, the animal was euthanized with a solution of pentobarbital after 1 h under anesthesia to provide sufficient time for the expression of c-Fos and perfused with 4% paraformaldehyde (PFA). The brain was extracted, immersion-fixed in 4% PFA overnight, and embedded in paraffin. The tissue blocks were sectioned (5 µm) to extract horizontal sections. The sections were incubated with the primary antibody (mouse anti-c-Fos, scbt sc-8047, 1:500 dilution) at 4°C overnight and then incubated in secondary antibody (goat anti-mouse Cruz Fluor 488 (scbt sc-533653, 1:50 dilution) for 2 h in the dark at room temperature. A single section was analyzed for each brain extracted. Sections at -1.5 mm DV were imaged under a Nikon epifluorescence inverted microscope using GFP lighting with 10× magnification (Fig. 3(d)). All imaging parameters were constant across all captured images (2048 × 2040 pixels, 1274 × 1274 µm^2^). The left hemisphere which was not exposed to rMTI stimulation was used as the control side. Cells were counted using the “analyze particles” feature of ImageJ software (NIH, Bethesda, USA) which automatically detected fluorescent cells based on size (set to > 3 µm)) and shape (set to 0.5-1). A cell was considered positive when the intensity of the cell staining was significantly higher than the surrounding background (i.e, when above an image-specific threshold). The same entire experiment was repeated for the animals that received stimulation from 1 solenoid (11 kHz) and 2 solenoids provided with the same frequency signal (11 kHz + 11 kHz).

**Fig. 3:**
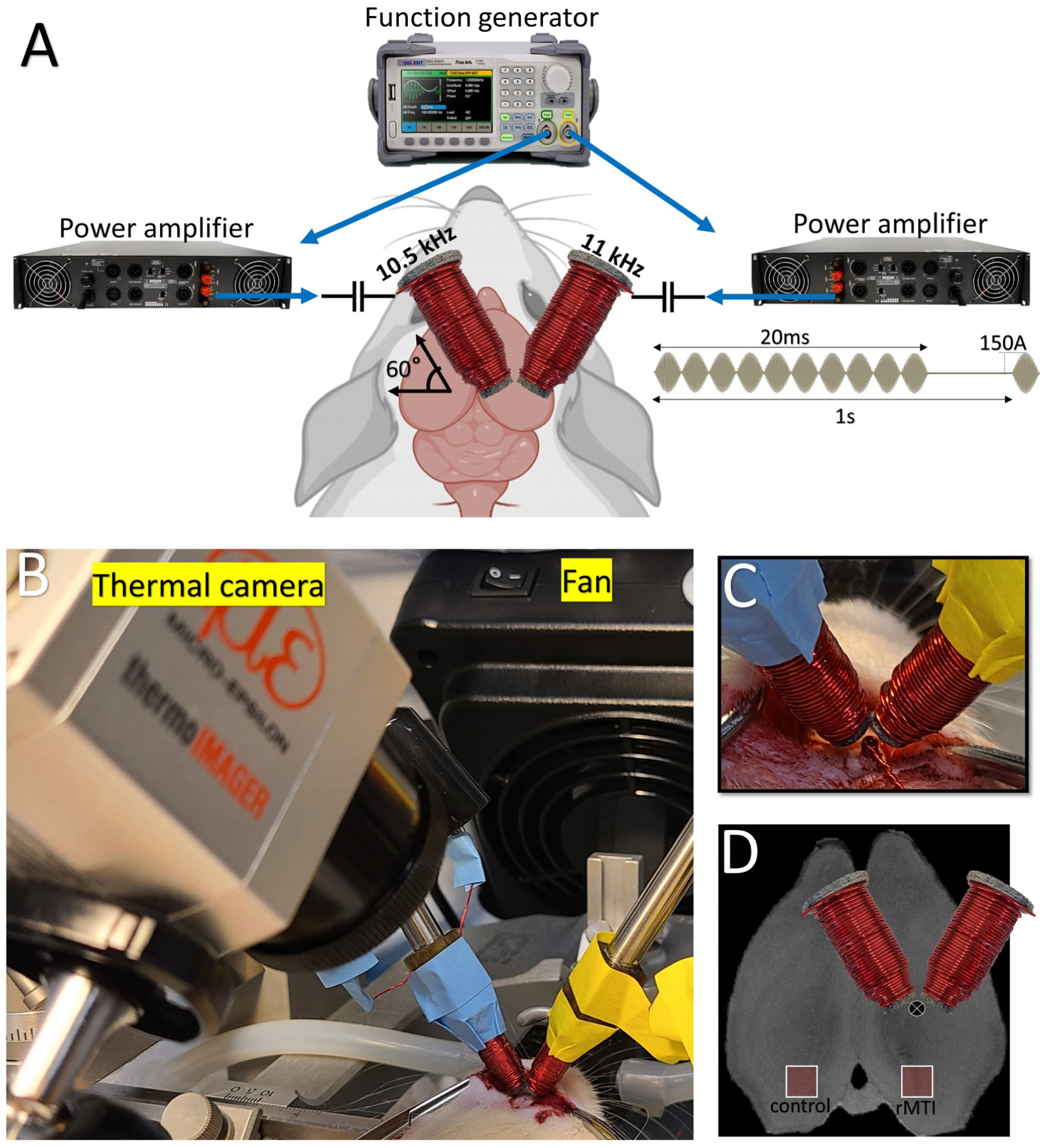
Images of the in-vivo experimental setup. (A) Drawing showing the instruments used during rMTI stimulation and the waveform delivered to the solenoids. (B) Picture of the animal experiment showing the fan, thermal camera, and tilted solenoids placed over the skull. (C) Close-view of the search coil glue onto the skull that was used to calibrate the delivered currents before rMTI stimulation. (D) Drawing showing the solenoid placement and the locations of imaged brain regions located in the visual cortex (represented as a red square).

## III. Results

### A. Single solenoid construction and characterization

The solenoid is shown in Fig. 4, and its parameters and specifications are summarized in Table I. Note that the L, R, and Q measurements were taken when the solenoid was connected to long cables (∼1.21 m) that were used to connect the solenoids to the power amplifier. We constructed a small solenoid with 80 turns of copper wire. A 26 AWG (∼0.4 mm) wire was winded on a 3D printed tapered bobbin. In an attempt to reduce wire resistance, we also experimented with Litz wires but realized they did not provide an advantage over standard wires for frequencies below 20 kHz. The bobbin diameter and length were set to match that of the magnetic core. Using the optimized parameters shown in Table I, the coil achieves a Q of 10.2 at 11 kHz. We fabricated other solenoids with much larger quality factors, but they also generated more heat and had larger diameters, and thus were considered inferior designs for the proof-of-concept experiment conducted in this research. Fig. 5 shows the results for the measured and simulated B_z_-field peak against supplied current to a vertically aligned solenoid. Measurement results are in good agreement with COMSOL simulations.

**Fig. 4:**
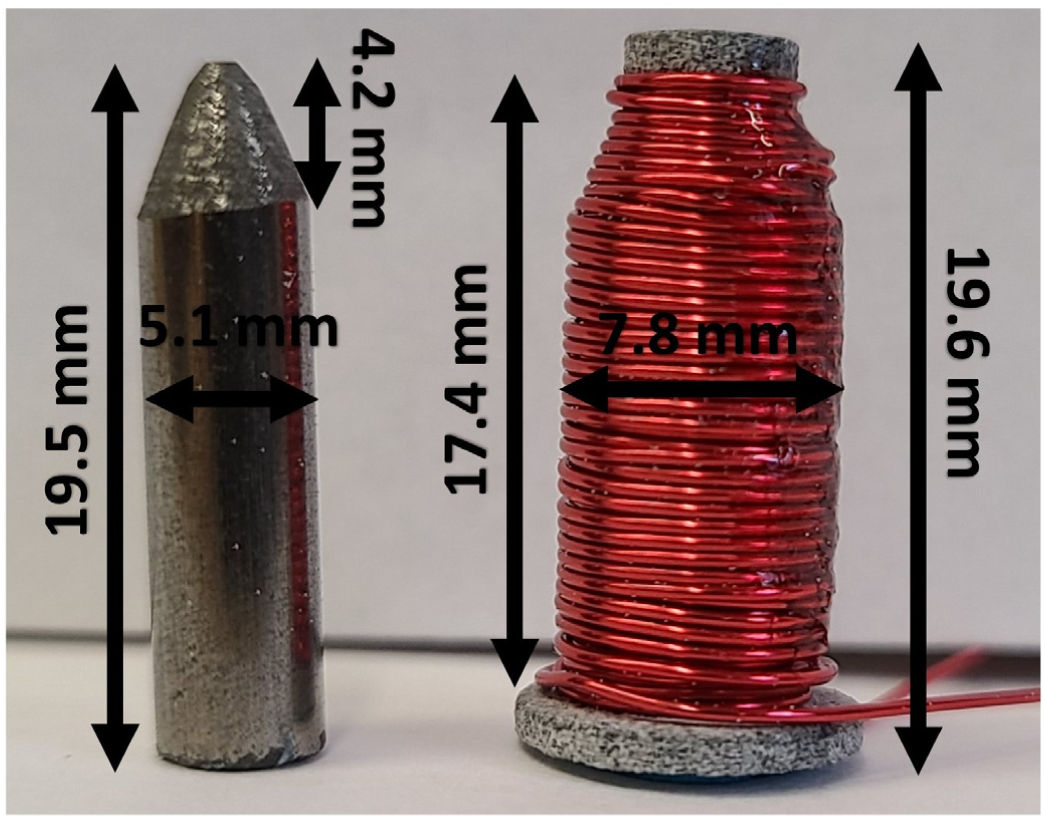
A photograph showing the novel solenoid and its dimensions. The tapered iron powder magnetic core (left) is inserted into the tapered bobbin (right).

**Fig. 5:**
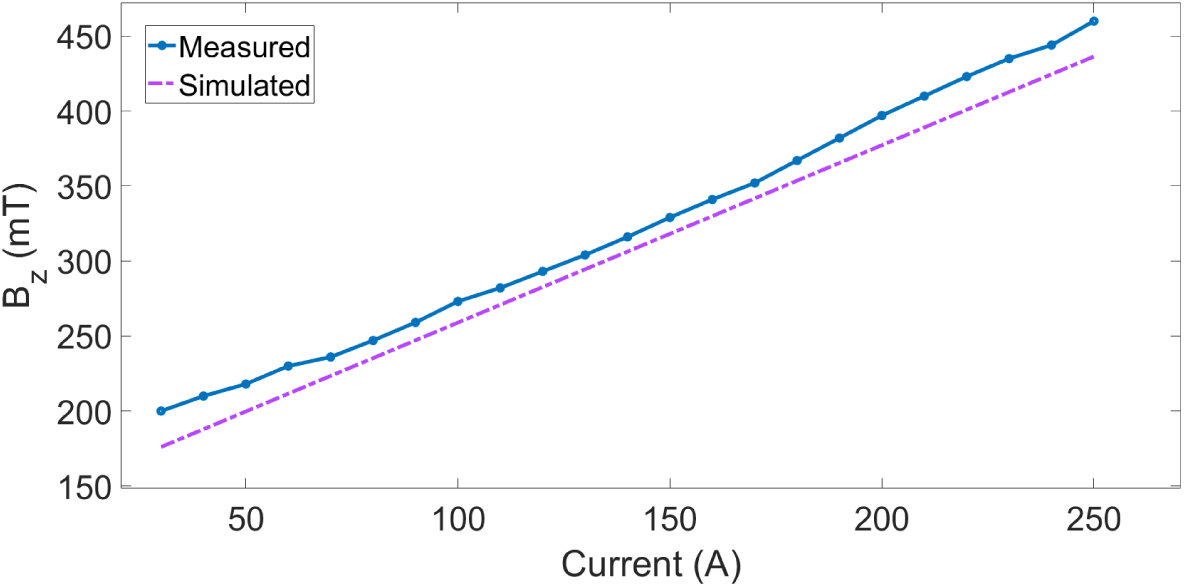
Measured and simulated B_z_-field peak against supplied current to one vertically aligned solenoid. The search probe was placed at 2 mm from the solenoid with a center-to-center alignment.

**TABLE I.** Solenoid Design Parameters and characterization

| Parameters | MTI Solenoid |
| --- | --- |
| Outer diameter (mm) | 7.8 |
| Length (mm) | 19.6 |
| Wire material and diameter (mm) | Cu, 0.4 |
| Number of turns | 80 |
| Core radius and length (mm) | 5.1, 19.5 |
| Core material | Iron powder |
| Core relative permeability | 9 |
| L (μH) at 11 kHz | 40 |
| R ($\Omega$) at 11 kHz | 0.27 |
| Q at 11 kHz | 10.2 |

### B. MTI characterization

Fig. 6 shows the simulated rMTI B_z_-field generated by 2 tilted solenoids. The figure also a comparison between measured with simulated B_z_-field strength as a function of time (shown for only 2 cycles) for different x, y locations. As expected, the maximum B_z_-field is found where the wire is closest to the XY plane. The plots show that the measured maximum B_z_-field is slightly higher than that of the simulations. The small mismatch could be caused by the distorted measured envelope. The measured signal is distorted for two main reasons. Firstly, the power amplifier (PA) does not output a perfectly linear signal since the solenoid is not matched to the output impedance of the PA. Secondly, although the solenoids are tilted to minimize coupling, they are still weakly magnetically coupled due to their proximity to each other. We do not expect the coupling to be this strong in a real application where the solenoids are separated by a much larger distance. The modulation depth (AM), which is a percentage that represents the amplitude variation of the envelope, varies considerably for the different x, y locations shown. For instance, the measured B_z_-field waveform at x = 0, y = 0 (location #1) shows an AM above 90%, whereas the measured B_z_-field waveform at x = 0, y = 1.35 mm (location #2) shows an AM below 60%. This signifies that the TI focal point is relatively small which enables focal rMTI stimulation. At x = 0, y = 5 mm (location #3), the AM is significantly attenuated as the beating envelope is barely visible. At x = 3.5, y = 0 mm (location #4), the AM is above 80%, signifying that the beating envelope along the x-axis does not attenuate as fast as on the y-axis.

**Fig. 6:**
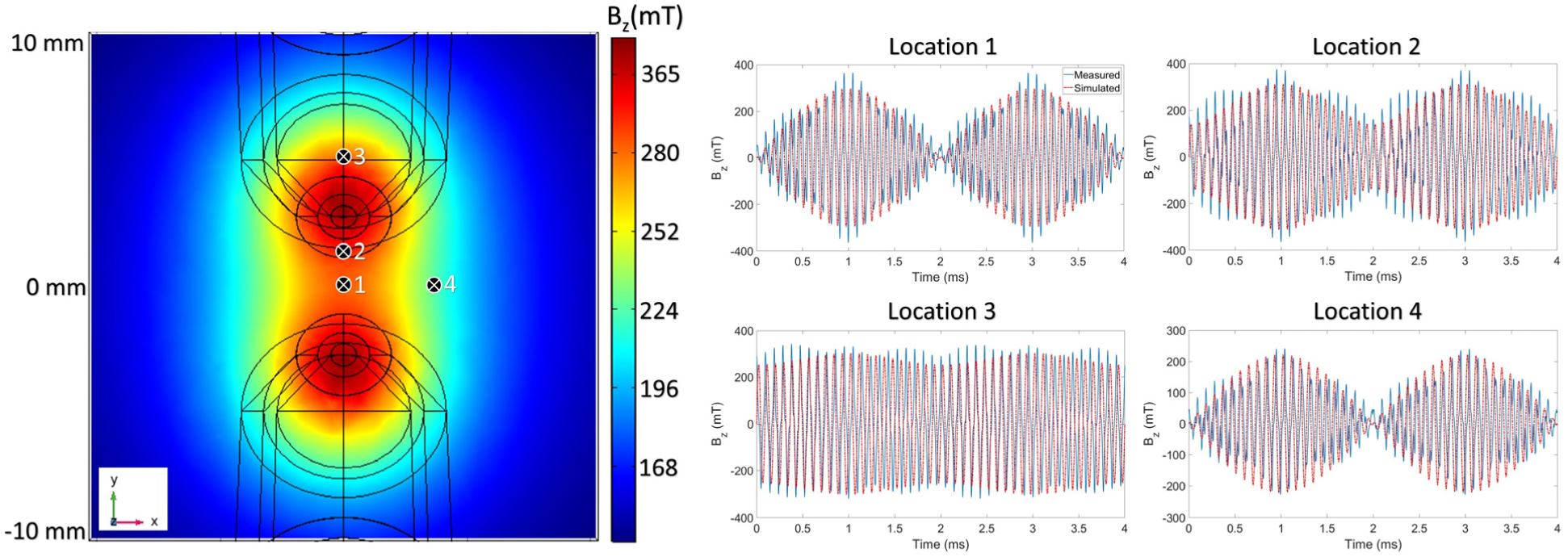
Measured and simulated rMTI B_z_-field generated by 2 tilted solenoids. (Left) Top-view showing the simulated B_z_-field distribution on a xy plane when 150 A was applied to both solenoids. One solenoid received a 10.5 kHz sine wave while the other received a 11 kHz sine wave. (Right) Comparison between measured and simulated rMTI B_z_-field against time shown at 4 locations. The 500 Hz envelope formed by the temporally interfering fields is plotted for a time length of 4 ms.

Fig. 7 shows the simulated rMTI E_env_-field generated by 2 tilted solenoids. The spatial distribution of E_env_-field shows two hot spots along the x-axis, thus indicating that we should not be expecting neuronal activation between the solenoids but about 5 mm below or above the center along the x-axis.

**Fig. 7:**
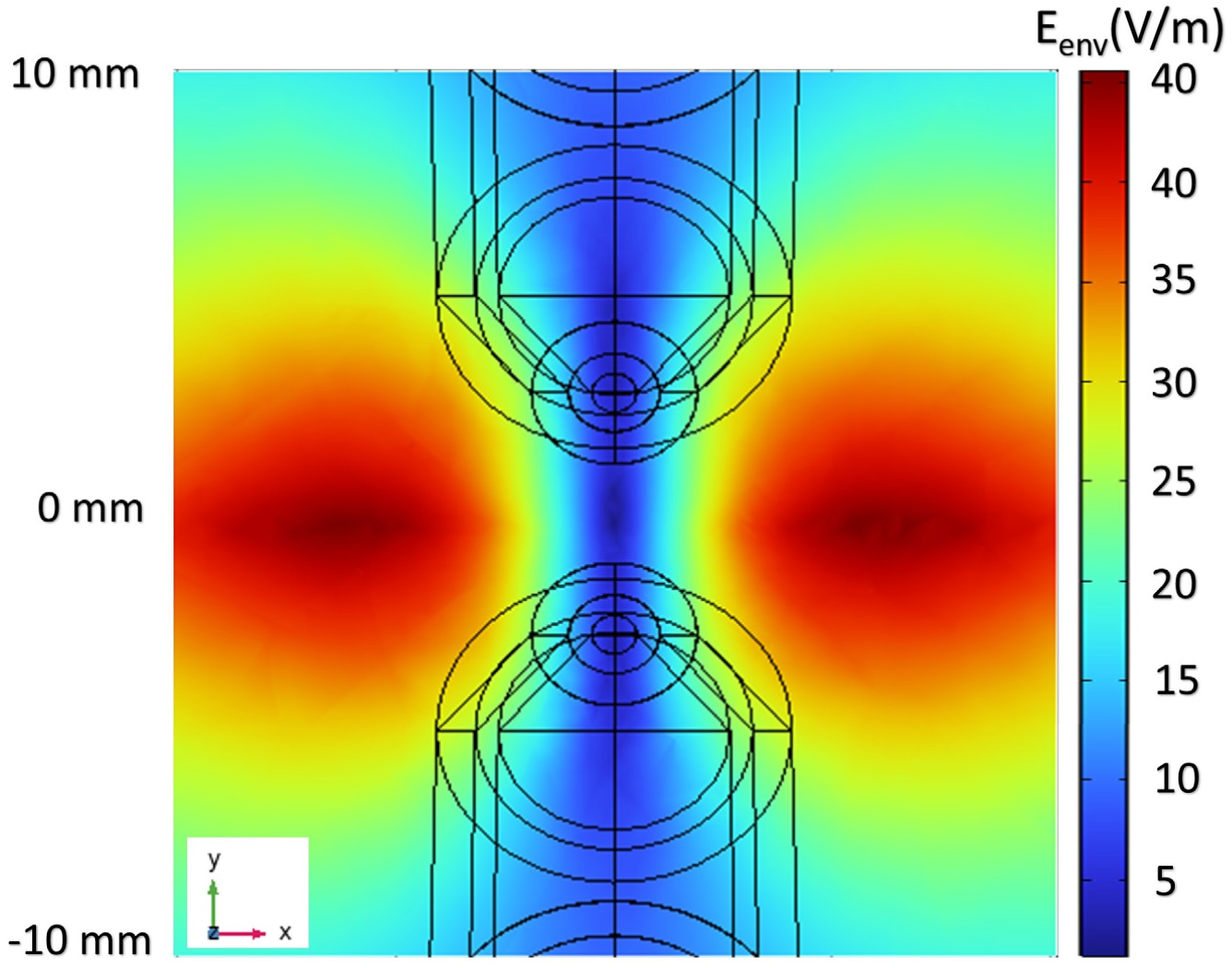
Spatial distribution of the envelope of the interfered electric fields generated by two solenoids.

The heat generated by the solenoids was imaged (Fig. 8(Top)) and plotted over time (Fig. 8(Bottom)). The temperature of the solenoid stops increasing after 6 min. After 15 min of rMTI stimulation (∼9900 cycles), the temperature of the solenoid increased by 40°C. This increase would have been much larger without the use of a fan (i.e, >60°C). The temperature at the center of the solenoid is about 8°C higher than that of the tip of the solenoid. The heat generated by the two solenoids caused the skull temperature to increase by 2.2°C which is below our safety limit of 2.5°C.

**Fig. 8:**
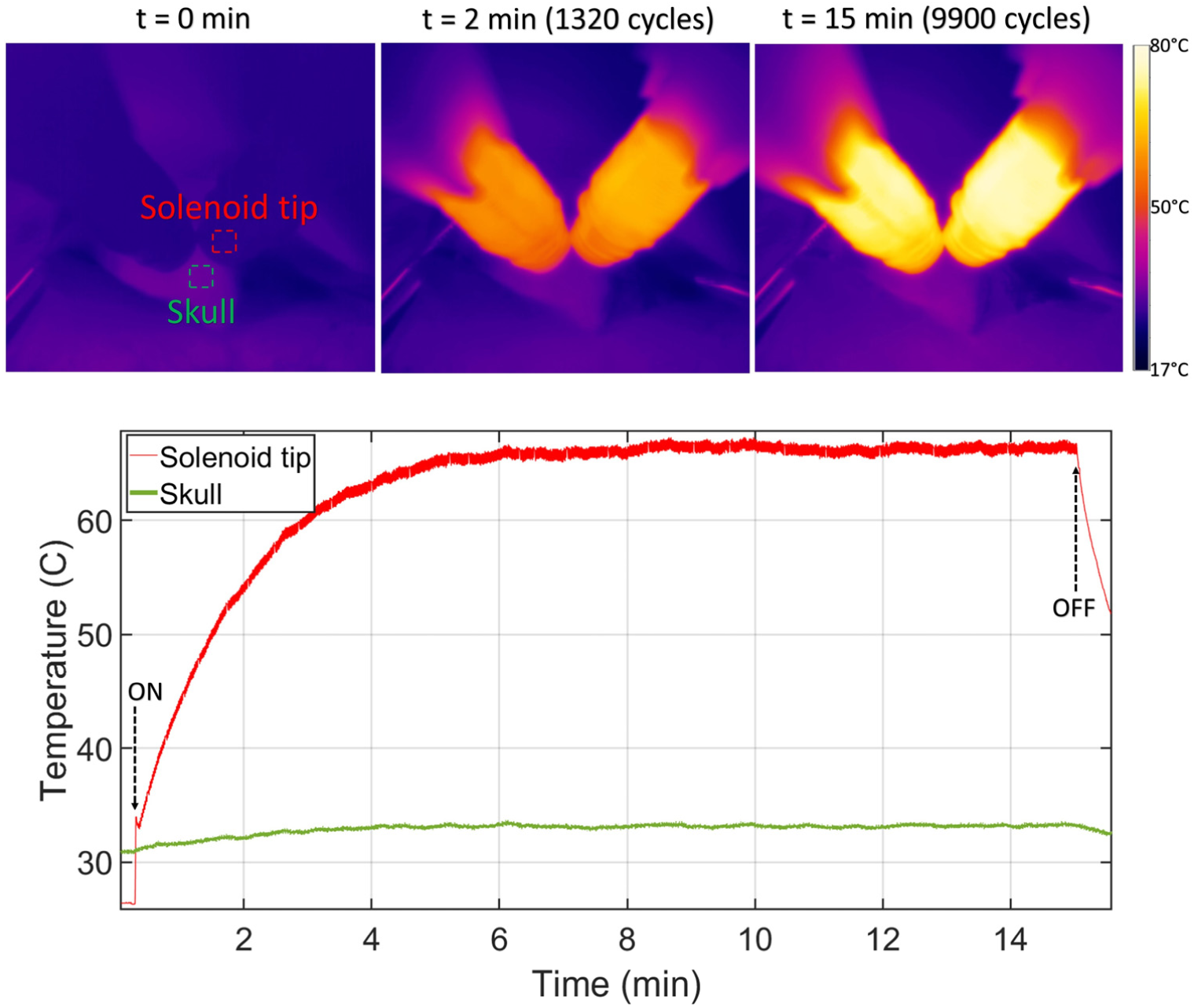
Temperature measurements taken using a thermal camera during rMTI stimulation at 150 A. (Top) Three pictures of the solenoids placed above the rat scalp taken at time 0, 2 and 15 min. (Bottom) Plot showing the temperature of the solenoid tip and rat scalp vs. time. The skull temperature increases by ∼2.2°C during rMTI stimulation.

### C. rMTI stimulation of a rat brain

Fig. 9(a) shows representative immunofluorescence images of each stimulation group showing c-Fos expression in sections extracted from the cortex of rats. In rats exposed to rMTI stimulation, we observed intense c-Fos activation in the visual cortex (more precisely, from bregma, AP -7 mm, ML -2.5 mm, DV -1.5 mm) which is below the solenoids’ target site coordinates. Therefore, as expected, there is up-regulation of c-Fos within 1 mm of the E_env_-field hot spot. As shown in Fig. 9(b), the number of c-Fos positive cells is more than 3 times higher in the right hemisphere, which received rMTI stimulation, compared to the left hemisphere which served as a non-stimulation control. Furthermore, in rats that received single solenoid stimulation or stimulation from solenoids supplied with the same sinusoidal frequency (11 kHz + 11 kHz), c-Fos expression showed similar levels as control, suggesting that the formation of a beating envelope is required to increase the number of c-Fos positive cells.

**Fig. 9:**
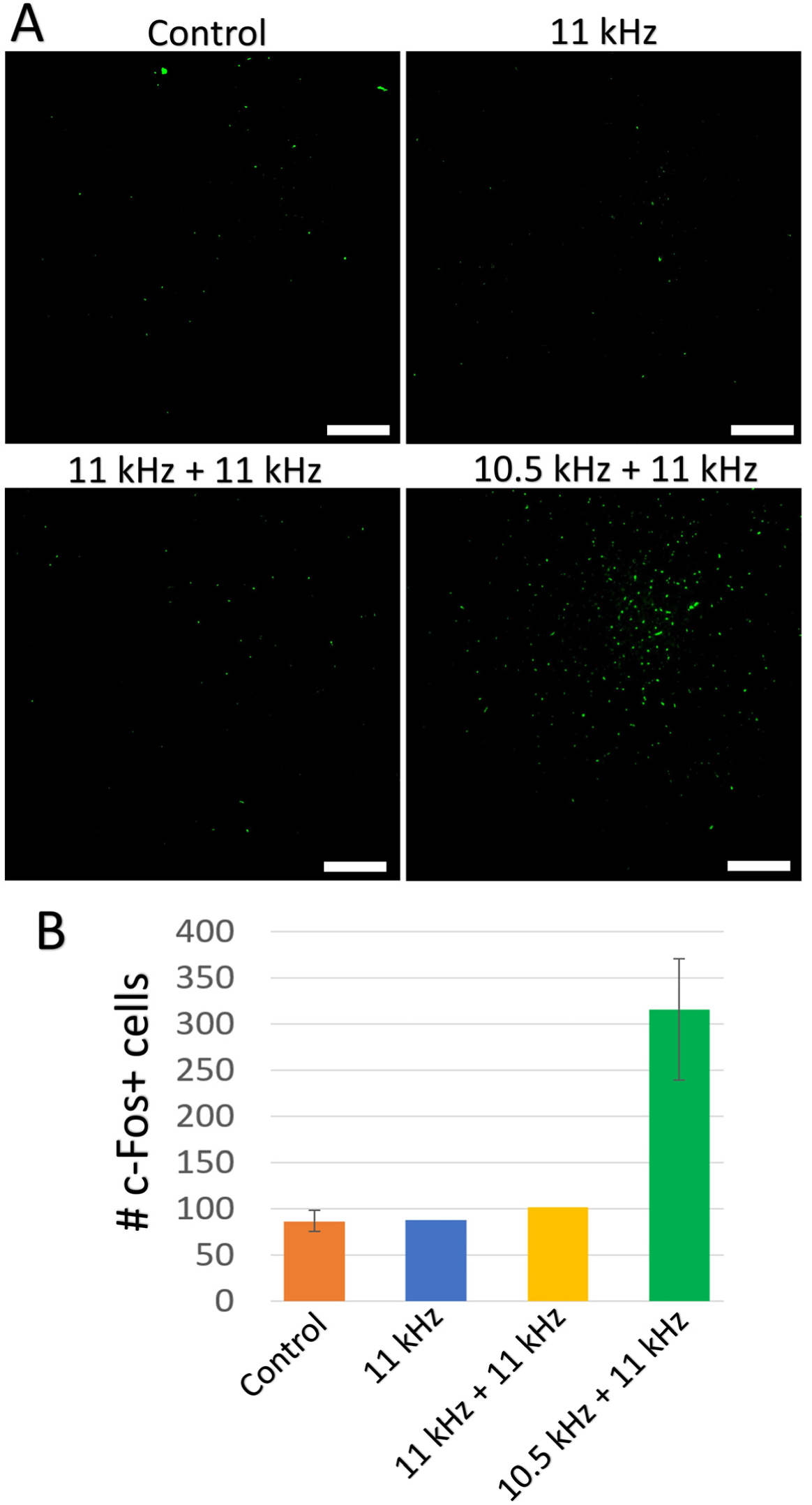
Representative immunofluorescence images of each stimulation group showing c-Fos expression in the visual cortex of rats. (A) Example images showing less expression of c-Fos in the non-stimulated side (control) and high frequency stimulation (without beating envelope) side vs. upregulation of c-Fos in the rMTI stimulated side. (B) Quantification of c-Fos expression within each image for control (n = 3), 11 kHz stimulation (n = 1), 11 kHz + 11kHz stimulation (n = 1), and 10.5 kHz + 11 kHz rMTI stimulation (n = 3). Scale bar is 200 µm.

## IV. Discussion

To investigate the effect of rMTI stimulation, we looked at neuronal c-Fos expression which is a member of the family of the immediate-early genes. In this work we did not record neuronal activity directly because of the difficulty of obtaining clean neurophysiology in the setting of continuous oscillatory magnetic fields. Although our experiments are preliminary and include only a small number of animals, our results demontrsate c-Fos elevation in brain regions targeted by rMTI stimulation, thus supporting the capability of rMTI to be used for non-invasive focal stimulation. Comparison with control stimulation conditions supports that the temporal interference is responsible for the elevation in c-Fos expression because stimulation induced by a single solenoid, and stimulation induced by two solenoids that were supplied with the same frequency signal did not appear to alter c-Fos expression (although only one animal was tested for each condition). A change in expression of c-Fos is generally considered to reflect a change in cellular activity, and in neurons is associated with increased electrical activity or a response to other factors (e.g., brain-derived neurotrophic factor expression). However, c-Fos expression has also been detected in non-neuronal cells. As a next step, therefore, it will be important to perform similar experiments but use either other, more specific, trackers of neuronal activity such as Npas4 or to perform double labelling experiments to identify the cell type that is activated by rMTI [28], [29].

To better utilize or improve rMTI stimulation technologies, we must first understand how the MTI E-field influences the transmembrane potential of neurons to induce activity or plasticity. There is controversy in the literature regarding the mechanism responsible for making neurons react to electrical temporal interference stimulation. On one hand, Grossman *et al*. [17] hypothesize that the low-pass filtering properties of neurons are the underlying mechanism for TI stimulation and that strong kHz-frequency electric fields do not (or minimally) lead to off-target tissue conduction blocks. On the other hand, Mirzakhalili *et al*. [30] claim based on mathematical modeling that TI stimulation activates neurons through nonlinear processes which are capable of “demodulating” the TI waveform and cause a conduction block in off-target tissues. Our understanding is different from that of these two groups. We hypothesize that the activity observed with TI stimulation is the result of “onset response” which typically precedes a conduction block [31]. As described by Niloy *et al*. [32], an onset response is an asynchronous bursting of neural activity before a block. We speculate that when the E-field beating envelope approaches 0 V/m, the membrane potential of neurons that are exposed to TI return to their resting membrane potential making the neurons more susceptible to another onset response shortly after being blocked. Therefore, each beating cycle may create a new onset response for neurons located within the focal point of TI, whereas off-target neurons manifest the onset response only once and remain blocked during the entire continuous waveform generated by rMTI stimulation.

A figure-of-eight TMS coil induces an E-field of about 150-250 V/m [8], [33]. The rodent-specific solenoid constructed for rMTI stimulation exhibits a maximum E-field of 40 V/m when 150 A is delivered to the solenoids. Therefore, we do not know for certain whether the rMTI E-fields are inducing subthreshold or suprathreshold stimulation. Although the E-field is much smaller than that generated by human TMS coils, the rMTI waveform was designed to induce repetitive stimulation within 4 ms since it is composed of 10 beating envelope cycles.

Miniaturization of solenoids/coils has proven to be a significant hurdle. For instance, it was shown to be very challenging to directly elicit motor evoked potentials with custom solenoids/coils for rodents [34], [35]. The primary limiting factor is heat generated by the solenoids. The induced electric field does not scale with the diameter of the solenoid [35], [36]. Consequently, to produce the same electric field in the rodent brain as in a human brain the same current is required. This is a major challenge as very large currents flowing through miniaturized coils/solenoids that have smaller wire diameters will generate significant heating. This challenge is even more pronounced for rMTI solenoids since the current is continuously applied for longer durations as opposed to conventional TMS waveforms (single pulse is < 1ms). In conventional TMS, the current pulse is provided by rapidly discharging a capacitor through the coil/solenoid, whereas in the case of rMTI stimulation, the current waveform is a sinusoidal wave provided by a power amplifier. When the heat generated by the solenoid/coil through Joule heating increases the scalp temperature by more than 5°C, then the animal starts to feel pain [37]. We thus decided to limit the scalp and skull heating to 2.5°C which provided a sufficient safety margin. Our temperature measurement show that the temperature of the skull increases by 2.3°C during rMTI stimulation when 150 A was provided to the solenoids. It is thus clear that the maximum generated B-field and E-field is set by the temperature increase of the skull (or scalp for non-invasive rMTI stimulation) and not the power amplifier which can provide significantly larger currents. Therefore, the temperature measurements show that 150 A is approximately the limit for the constructed novel rodent specific solenoids. The stringent current limit makes it very challenging to target deep brain regions with the miniaturized solenoids.

Since we decided to stimulate cortical areas instead of deep brain structures, the solenoids have to be close to each other to enable focal rMTI stimulation. Both solenoids were tilted such that they formed a 60° angle. We have done additional simulations that shows that increasing the solenoid angle above 60° leads to much larger generated electric fields. However, our simulations did not consider the coupling between the solenoids. Therefore, in practice, tilting the solenoids with angles above 60° caused more distortion to the sinusoidal signals which in turn distorted the temporal interference. Knowing that there is a tradeoff between electric field intensity and envelope modulation depth, we found that a 60° angle was optimal.

Nonetheless, the rodent-specific solenoids used here were relatively easy to fabricate, and it would not be challenging to modify the human figure-of-eight TMS coils for rMTI stimulation. The present results are encouraging but future studies will need to address the mechanisms responsible for rMTI and its impact on neural activity. Most importantly, the question of whether a sodium depolarization block is occurring in off-target regions needs to be answered. The next iteration of this work will involve rMTI stimulation in larger awake animal models which allows for the construction of larger solenoids that produce less heat and are theoretically capable of stimulating deep regions in the brain without activating the overlying tissues.

## V. Conclusion

A new generation of TMS systems that allows for stimulation beyond the resolution and depth of “figure-of-eight” coils will give researchers more opportunities to target specific neural circuits for the treatment of neurological and neuropsychiatric diseases. We have proposed a novel brain magnetic stimulation technique that was inspired by previous works based on temporal interference. rMTI stimulation has the potential to become a clinical tool that can be used to directly target malfunctioning circuits deep in the brain, ultimately providing opportunities for better treatment of neural disorders in humans. In a step towards achieving this goal, we focused on developing novel solenoids for rMTI stimulation capable of focal neuronal activation in a rat model. We encourage further research studies on the mechanism of TI stimulation as better understanding the concept will lead to better technologies and solutions to challenges.

